# A scoping study of the whole-cell imaging literature: a foundational corpus, potential for mesoscale data synthesis, and implications for standardization of an emerging field

**DOI:** 10.1101/2025.02.03.636363

**Authors:** Mary Mirvis, Brooke Weingard, Steven N. Goodman, Wallace F. Marshall

**Affiliations:** Department of Biochemistry & Biophysics, University of California, San Francisco, San Francisco, CA; Department of Epidemiology and Population Health, Stanford University, Stanford, CA

## Abstract

The level of cellular organization bridging the mesoscale and whole-cell scale is coming into focus as a new frontier in cell biology. Great progress has been made in unraveling the complex physical and functional interconnectivity of organelles, but how the entire organelle network is spatially arranged within the cytoplasm is only beginning to be explored. Drawing on cross-disciplinary research synthesis methods, we systematically curated the whole-cell volumetric imaging literature, resulting in a corpus consisting of 89 studies and 118 image datasets. We describe the trajectory and current state of the field between 2004 and 2024. A broad characterization, or “scoping review”, of bibliometrics, study design, and reporting practices shows accelerating technological development and research output. We find high variability in study design and reporting practices, including imaging modality, model organism, cellular contexts, organelles imaged, and analyses. Due to the laborious, low-throughput nature of most volumetric imaging methods, we find trends toward small sample sizes (<10 cells) and small cell types. We describe common quantitative analyses across studies, including volumetric ratios of organelles and inter-organelle contact analyses. This work establishes the initial iteration of a growing dataset of whole-cell imaging literature and data, and motivates a call for standardized whole-cell imaging study design, reporting, and data sharing practices in the context of an emerging sub-field of cell biology. Our curated dataset now provides the basis for a plethora of future aggregate and comparative analyses to reveal larger patterns and generalized hypotheses about the systems behavior and regulation of whole-cell organelle networks. More broadly, we showcase the potential of new rigorous secondary research methods to strengthen cell biology’s literature review and reproducibility toolkit, create new avenues for discovery, and promote open research practices that support secondary data-reuse and integration.

## INTRODUCTION

Biological organization spans many physical scales, from the atomic and molecular, to global ecosystems. In cell biology, the dominant relevant scale is that of the cell as the indivisible unit of life, where questions of cellular function, behavior, motion, communication, and shape dominate. At the subcellular scale of molecular biology and biochemistry, the structure, function, information flow of biological molecules, from ions to large macromolecular complexes such as nucleic acids and ribosomes reigns. Between these two scales lies the less clearly defined mesoscale, encompassing all subcellular structures larger than ∼100 nm including condensates, cytoskeletal assemblies, organelles, and the cytoplasm (Sear et al., 2015). Great progress has been made in identifying the relevant structural components of the mesoscale, but defining the driving forces and principles governing the complex spatial patterning, dynamics, interactions and systems properties of mesoscale entities is now a pressing challenge in cell biology.

At this frontier of biological organization, new types of questions arise at the intersection of cell, systems, and physical biology. How do well-understood molecular mechanisms scale *en masse* across the entirety of the cell? How do disparate mesoscale structures, all well known to be highly dynamic and structurally complex, interact with each other? To what extent are mesoscale structure and function directly governed by genetically programmed or energetically active mechanisms versus emerging or self-organizing from intrinsic mechanical interactions, self-organization (Laughlin et al., 2000)? Can the entire set of mesoscale components be understood as a whole interconnected system, and what conceptual frameworks and experimental and theoretical approaches are necessary for this understanding?

Organelles, the largest mesoscale structural components, together represent the highest level of organization within the mesoscale. The fine ultrastructure of organelles has been extensively characterized through electron and light microscopy since the 1950s, and a rich understanding of the complex molecular composition, cellular functions, and dynamics of each individual organelle has been amassed since the 1980s. In the last decade, a new picture of organelles as interconnected components in a complex and dynamic network mediated by many contact site proteins and communication mechanisms has emerged, summarized in many reviews (Cohen et al., 2018; Scorrano et al., 2019; Voeltz et al., 2024). A systems view of organelles spawns new questions around the structure, dynamics, and regulation of the organelle network; its responsiveness to perturbations and mechanism of disruption in disease; and the role of each organelle component and pairwise interaction in network structure and function. Questions of this nature are being increasingly addressed through systems imaging (Bray et al., 2016; Rohban et al., 2017; Wang & Mukherji, 2022), spatial proteomics (Cho et al., 2022; Hein et al., 2023; Lundberg & Borner, 2019; Qin et al., 2021), and computational modeling (Donovan-Maiye et al., 2022; Earnest et al., 2017; Majarian et al., 2019; Murphy, 2012; Viana et al., 2023; Zhao & Murphy, 2007).

### A new construct: Cell Anatomy

Studying the structure of organelle networks is a challenging task. Highly dynamic and morphologically heterogeneous organelles interact physically through a combination of protein-based membrane contact sites (MCS), mechanical forces, spatial constraints, membrane dynamics, and potentially other unidentified mechanisms. In aggregate, the coalescence of organelle morphologies, spatial distributions, and inter-relationships produces whole-cell patterns, which we term “cell anatomy”. Cell anatomy is a fundamental, but still incompletely defined cellular property bridging the mesoscale and whole cell, which is presumably both unique to specific cell types, as well as variable and dynamic within and across cell states. By defining and observing cell anatomy across many cells and contexts, we can formulate novel questions. Which types, features, or variations of cell anatomy are common, and which are context-specific? Which are genetically regulated versus determined by simple geometry or intrinsic physics? Answering such questions requires the generation of microscopy images of multiple organelles in tandem in the context of the entire cellular volume, revealing the morphology and positioning of each organelle in its native, three-dimensional context.

Whole-cell imaging has been accomplished across a range of cell types using a variety of imaging modalities, including electron, X-ray, and light microscopy methods. Among image data types, whole-cell imaging is particularly technically challenging. Images encompassing large biological volumes[1] must be acquired and reconstructed at resolutions high enough to accurately capture and segment multiple subcellular structures. Segmentation must be done either manually, which requires specialist expertise and many hours of labor per image, or automatically, which requires sufficient amounts of high-quality manually-segmented training data and rigorous validation (Chen et al., 2024; Ekman et al., 2020; Erozan et al., 2024). Addressing questions about the variability and regulation of 3D mesoscale patterns requires data from large numbers of cells and is therefore generally beyond the scope of single whole-cell imaging studies. Aggregating the existing set of relatively small whole-cell imaging studies may reveal trends in cell anatomy across cell types and biological contexts, providing an avenue for novel analyses and insights into cell anatomy and organelle interaction networks.

This goal requires finding and curating relevant whole-cell imaging studies across the literature as comprehensively as possible, extracting useful information, and analyzing image data and reported measurements. Under the broad umbrella of secondary research, there exists a wide array of methods to synthesize and review published work, yielding summaries of existing knowledge, meta-scientific insights into the state of a field or subset of literature, and identification of trends and gaps in the research landscape to motivate future work. A large family of systematic, rigorous, transparent, reproducible approaches to literature review collectively known as *evidence synthesis* or *research synthesis*, offers an array of rigorously developed methods for comprehensively and reproducibly curating and analyzing trends and knowledge across studies (Munn et al., 2018, 2023; Tricco et al., 2016). Here, we present a first step towards leveraging these tools for synthesizing mesoscale data from whole-cell imaging studies.

Our evidence synthesis approach ultimately seeks to uncover general and context-specific trends in cell anatomical patterns and organelle interdependencies. Combining information from many publications and datasets is a promising avenue to answer larger-scale questions than could be addressed by the results of individual published works. In order to formulate specific testable hypotheses, it is necessary to first have a clear picture of the amount and content of data currently available, and the ways it can be merged and compared across studies. Therefore, a crucial first step is to produce a thorough characterization of the volume, quality, and technical and biological features of the whole-cell image data landscape. In this work, we describe a “scoping study”, based on “scoping review” methods previously described and standardized (Arksey & O’Malley, 2005; Tricco et al., 2018). We present the development of our scoping methodology, including iterative search strategy development and screening, and delineate the contours of the whole-cell imaging field. We summarize publication trends and the range of methodologies, reporting practices, and biological conditions included in the rapidly growing corpus of whole-cell imaging data. We highlight key points of overlap in the data which lay the foundation for future systematic review work. Our work produces a foundational corpus of whole-cell imaging data with rich potential for comprehensive knowledge synthesis, novel biological insights, and a bird’s eye view of the changing trends and needs of this promising and rapidly-crystallizing sub-field of mesoscale cell biology.

## WORKFLOW

We adapted our approach from the well-defined scoping review methodology (Arksey & O’Malley, 2005; Peters et al., 2022). Our workflow for this study is summarized in Figure 1, and full details into each step of the workflow, including screening criteria, categorization, and analysis, are described in Methods. In brief, we developed a search strategy consisting of a set of microscopy terms and a set of cell- and organelle-scale terms, based on an initial set of 11 target papers. Our search strategy returned over 11,000 results from PubMed. We defined two main inclusion criteria: 1) generation of original three-dimensional *whole-cell* image data in eukaryotic cells, including complete cell boundary delineation, and 2) volumetric information for at least two *organelles* concurrently must be included in the whole-cell volume. After three screening stages – title and PubMed summary, abstract, and full text – conducted by a team of three reviewers, 89 studies were fully included. We extracted information from studies to summarize publication trends and study design, including imaging modality and resolution, cell types, sample sizes, organelles, comparative conditions, and quantitative analyses

**Figure 1.**
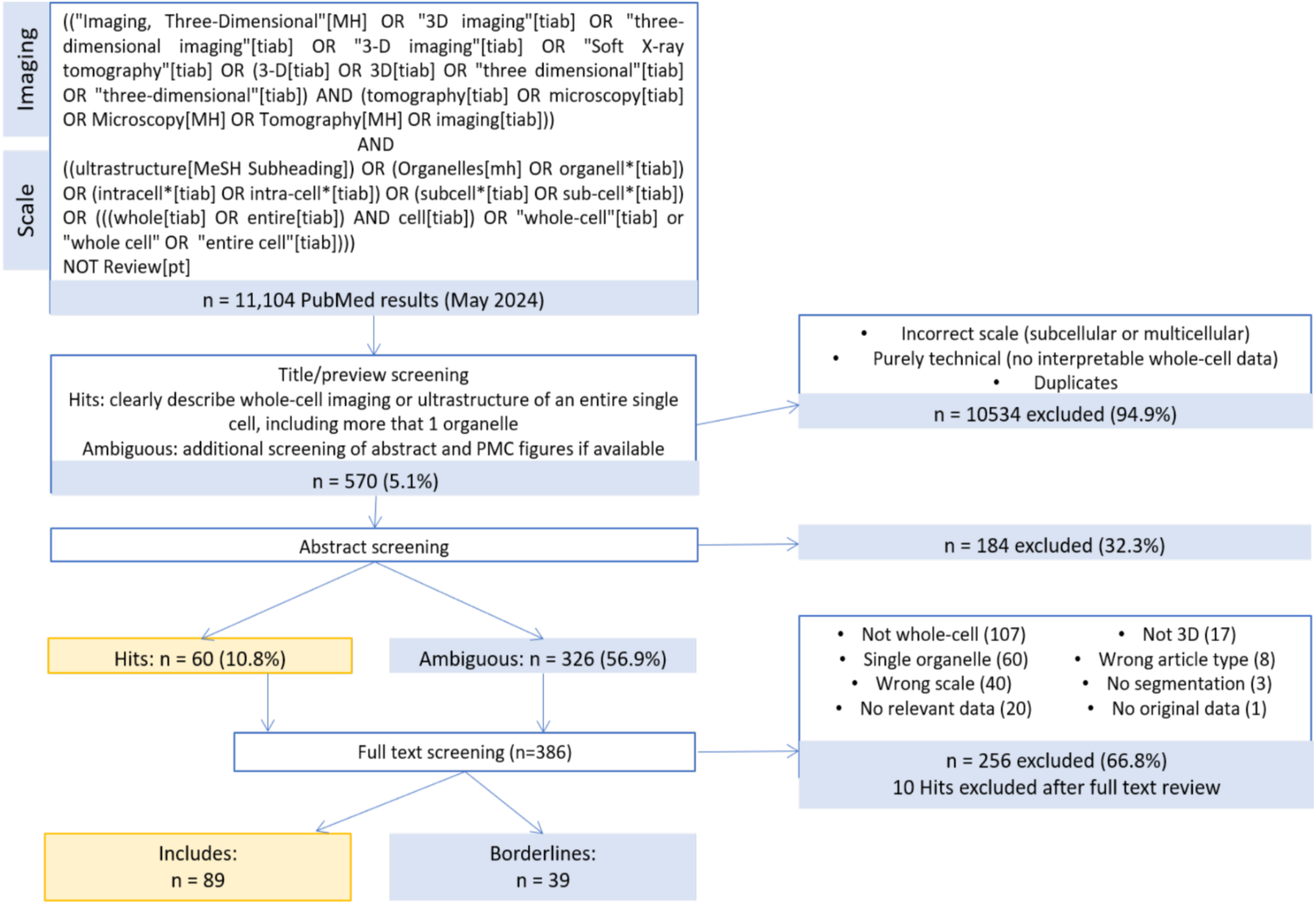
Study workflow to curate whole-cell imaging literature. PRISMA diagram detailing the PubMed search strategy and screening steps used, with included and excluded study counts at each stage. In search strategy, asterisk (*) indicates wildcard character, [MH] indicates Mesh header, [tiab] indicates title/abstract field, [pt] indicates publication type.

## RESULTS

### Metadata and bibliometrics

The potential for effective study curation and data extraction, and ultimately, novel insights into whole-cell structure, depends on the overall volume, accessibility, and type of studies screened. We first aimed to broadly characterize the publishing trends within our corpus. The publication rate of studies containing whole-cell imaging data has risen steadily since 2005, with a peak at 14 papers in 2023 (the most recent complete year, as our search stopped in May 2024), and 49.4% of the included studies (44/89) published in the last 5 years (2019-2024) (Figure 2A). The majority are open-access publications, which is important for study amenability to systematic search and screening approaches (70/89, 78.7%) (Figure 2B).

**Figure 2.**
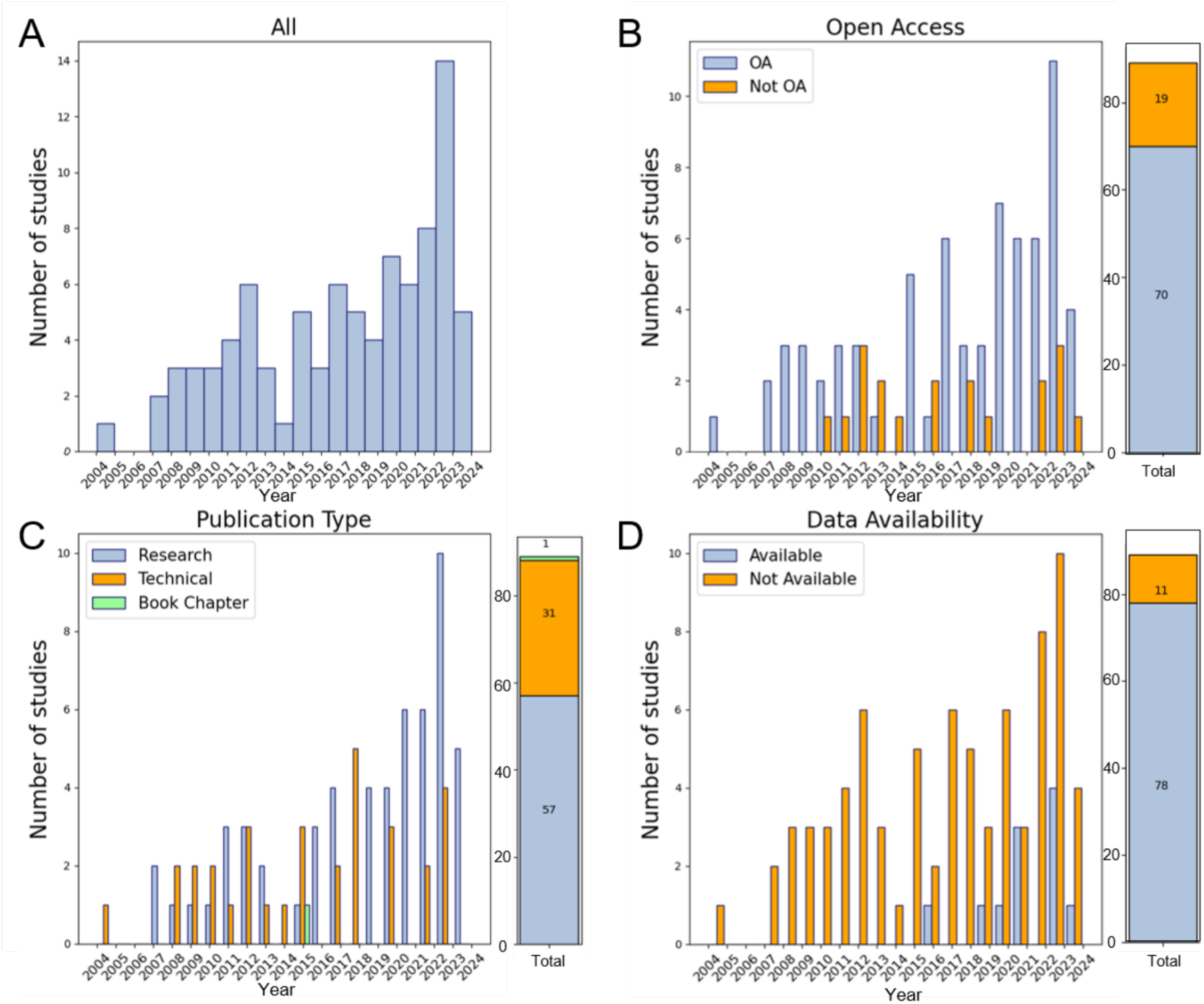
Scientometric summary of the whole-cell imaging literature. A) Histogram of total publication frequency by year. B) Open access (OA) publishing frequency by year. C) Publication type by year. D) Data availability, as reported, by year. Data are represented between 2004 and May 2024.

We reasoned that, while many of the studies we include may be similar in nature to our initial set, in that they perform extensive data collection and analysis in pursuit of a particular biological question, whole-cell image data may be found in other publication types. Technical studies which describe some advance in microscopy, image processing, or analysis, often provide small proof-of-concept datasets. While such studies may or may not undertake extensive biologically-driven analysis, the images themselves may include valuable structural information and be available for further analysis in combination with other datasets. The majority (57, 64%) were “research” articles, focused on answering a biological question, while 31 articles were considered “technical”, focused on method development, although some biological observations may be included (Figure 2C). One publication is a technical book chapter, which would have been otherwise excluded from our dataset as a secondary source, but included some original image data (Gluenz E et al., 2015).

We asked whether authors share the original imaging data via data repositories or other open-access platforms, which would enable secondary analysis of this information-rich data. Across the core included datasets and borderlines, only 11 included studies (12.4%) and 1 borderline study reported that underlying whole-cell image datasets were publicly available, although those that did were more recently published, from 2015 onwards (Figure 2D). Of these, seven datasets are available in their entirety as reported in the study (Cortese et al., 2020; Ezzedine et al., 2023; Laundon et al., 2019; Mocaer K et al., 2023; Müller et al., 2021; Rudlaff et al., 2020; Uwizeye et al., 2021a), while five are available in partial form (Antao NV et al., 2023; Hair et al., 2024; McQuate A et al., 2023; Sakaguchi M et al., 2016; Uwizeye C et al., 2021). A summary of publicly available datasets is provided in Additional File 3. Overall, we see a progressive expansion of study and data volume, as well as open science practices, with a pronounced increase in all of these trends in the past 5 years, suggesting good current and future potential for meta-scientific studies and secondary re-analysis of whole-cell imaging datasets.

Can whole-cell imaging represent an emerging field? We asked whether the corpus is scientifically interconnected, showing a substantial degree of scientific collaboration and scholarly acknowledgement. Analysis of co-authorship networks shows many tightly-linked co-authorship clusters (Figure S1A), and one large cluster of collaborators among prolific whole-cell microscopy researchers (authors on 2+ studies) (Figure S1B). A citation network shows that most (71/89), but not all studies in the corpus cite each other. A few studies (Larabell & Le Gros, 2004; Loconte et al., 2021; Uchida et al., 2009, 2011; Uwizeye et al., 2021b) emerged as the central, most highly connected studies. The most cited study (Valm et al., 2017), with over 1000 citations, does not cite previous studies in this corpus, but is cited by many subsequent papers that used other microscopy methods, indicating that there has been a convergence of whole-cell imaging efforts from disparate research streams. This bibliometric overview illustrates the growing pace and cohesion of the whole-cell imaging literature.

**Figure S1.**
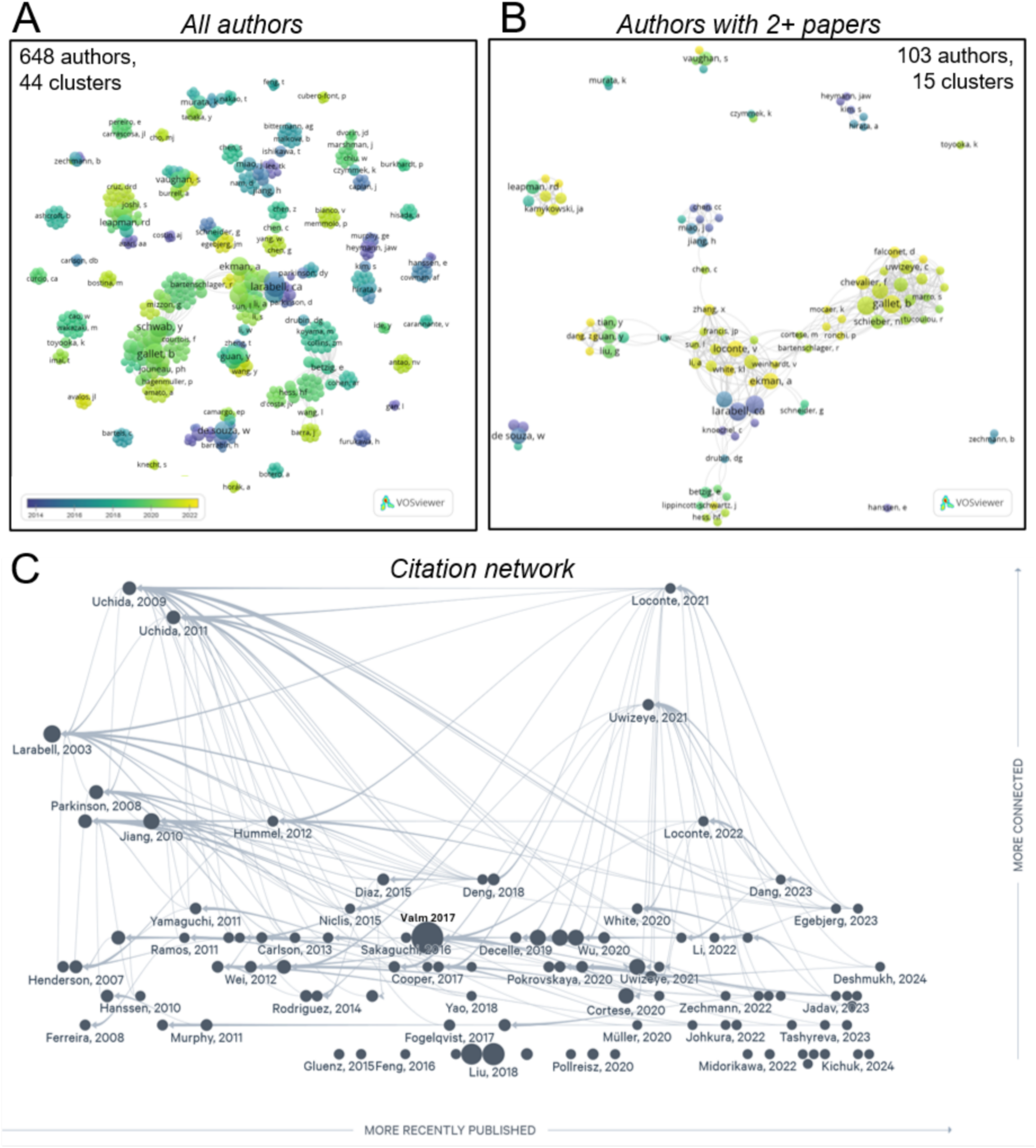
Bibliometric network analysis. A-B) Co-authorship network consisting of authors with A) any number of publications and B) those with at least two publications represented in the corpus. Color gradient represents average publication year per author and bubble size indicates relative number of articles per author. Visualizations created in VOSViewer. C) Citation network, created in Litmaps, representing most connected papers at the top, and most recent papers on the right. Bubble size reflects relative citation number, and arrows point to articles cited.

### Imaging modality and resolution

Structural biology through volumetric imaging has made great strides in recent years. The rapid improvement of image resolution, volumetric and statistical throughput, structure identification, and integration of cross-modal data, including correlative and dynamics approaches, have enabled exciting advancements in the overall accessibility and applicability of many imaging modalities to a wider and more complex array of problems than ever before. Whole-cell imaging approaches the upper limit of capabilities of most high resolution, 3D microscopy and reconstruction techniques. Each technique offers a slightly different set of strengths in terms of resolution, throughput, accessibility, cell type amenability and ultrastructural preservation, organelle identity and number, potential for quantitative analysis, and so on. Therefore, in characterizing the whole-cell imaging corpus, it is crucial to understand which imaging methods are being used, which are more common, and how they data they produce compare and contrast. This may motivate recommendations for modalities best suited for future whole-cell imaging studies.

In total, we identified 13 distinct volumetric imaging types of modalities used to produce whole-cell image data. Focused ion beam SEM (FIB-SEM) (also called ion-abrasion SEM, IA-SEM) and soft X-ray tomography (SXT) and are the most common modalities, represented in 22 and 21 studies, respectively (Figure 3A). These were followed by serial block-face SEM (SBF-SEM), serial sectioning TEM (ssTEM), electron tomography (ET), fluorescence (including 3D SIM, and multispectral lattice light sheet), optical tomography (including optical diffraction and optical computed tomography/live cell CT), hard X-ray tomography (HXT, X-ray ptychography), scanning TEM (STEM), and X-ray diffraction (XRD) techniques. Field emission SEM (FE-SEM), scanning transmission X-ray microscopy (STXM), and SEM array tomography (SEM-AT) were each found in one study. In total, 55 studies used electron microscopy (EM) techniques (FIB-SEM, SBF-SEM, ssTEM, ET, STEM, FE-SEM, SEM-AT), 27 (30%) used X-ray techniques (SXT, HXT, XRD, STXM), and 10 used optical techniques (fluorescence, optical tomography).

**Figure 3.**
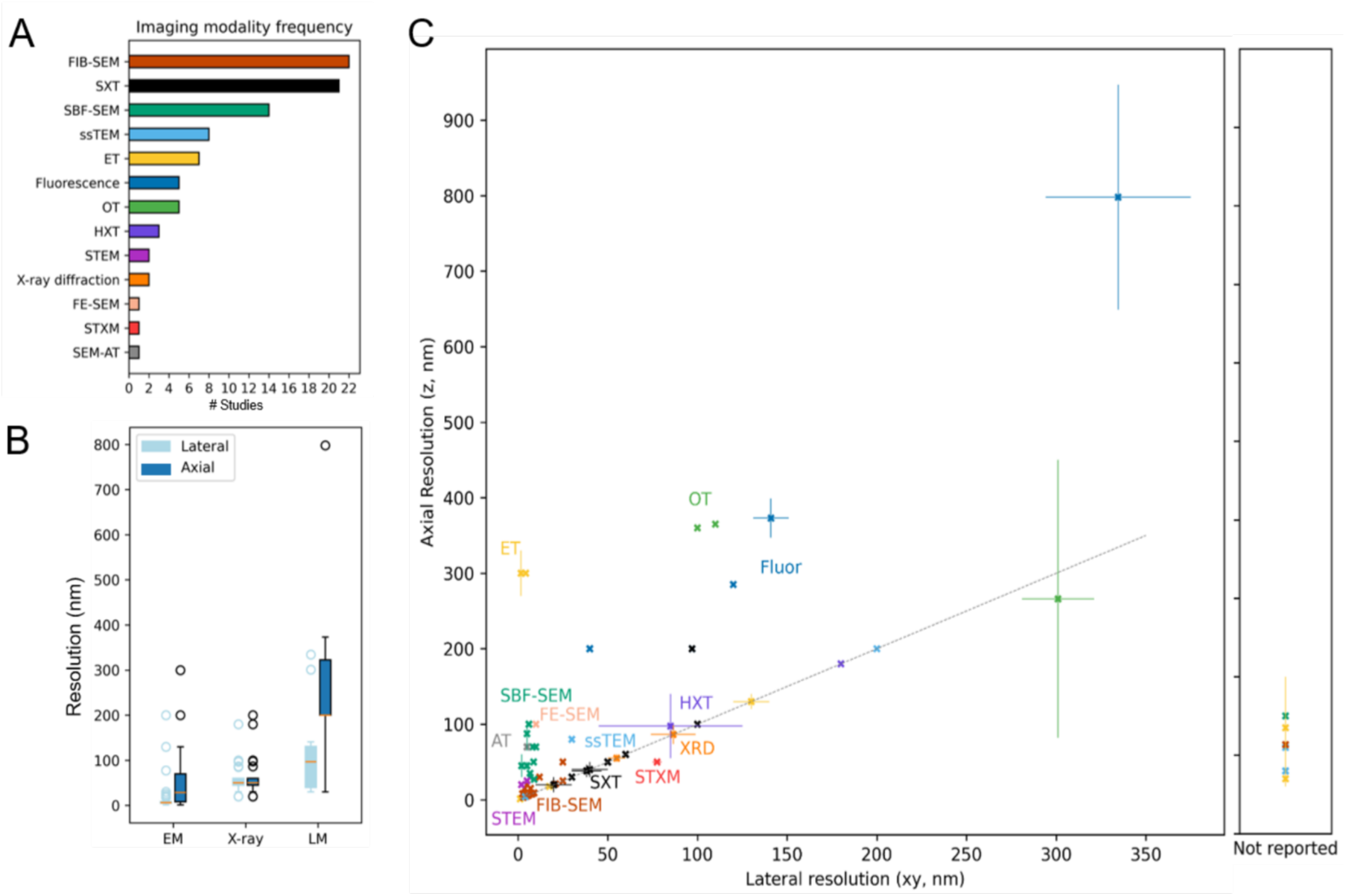
Summary of imaging methodology and resolution. A) The number of studies using each of the 13 imaging modalities. B) Box-and-whisker plots of reported lateral and axial imaging resolutions, in nanometers (nm), grouped by modality type: electron microscopy (EM), X-ray, or light microscopy (LM). A single value was plotted for each dataset. If multiple values or a range of values were reported, the value plotted is the median of the reported values. C) Scatterplot of axial vs lateral resolution in nanometers for each dataset, where reported, colored by modality. Error bars represent the reported range, when applicable, with the datapoint representing the median. The dotted line represents isotropic resolution (axial = lateral). Right panel represents datasets for which only axial resolution was reported.

Each modality excels within a limited range of spatial resolution, determining the size scale, volumetric completeness, throughput, and potential for artifacts of anisotropic resolution in the resulting images. Generally, optical imaging methods are limited by the wavelength of light (∼200nm), typically capable of achieving higher lateral (xy) than axial (z) resolution, but can be used to produce sample sizes in the hundreds or thousands. Electron microscopy modalities are capable of achieving resolution in the sub-10nm range, but are typically limited in volumetric scope and statistical throughput due to the high information density and cumbersome sample preparation, imaging, and image processing. X-ray based methodologies typically fall somewhere in between light and electron microscopy in throughput and resolution (Guo & Larabell, 2019; McDermott et al., 2012). The range of cell types potentially accessible for whole-cell reconstruction, information content contained in the resulting images, comparability across datasets and modalities, and capacity for secondary analysis is strongly dependent on image resolution. Whole-cell imaging requires the acquisition of large volumes, so even the best resolution possible with a particular modality or microscopy may need to be compromised to increase the reconstruction volume and/or throughput.

We extracted the technical parameter information for each distinct dataset across the corpus, where a dataset is considered a distinct set of cells of the same type imaged with the same modality, such that multiple datasets may appear within one study (e.g. four datasets comprising two cell types (*C. pepo, N. tabacum*) each imaged by two modalities each (ssTEM, FIB-SEM) in (Zechmann et al., 2022). We found 118 total datasets across the 89 studies (Additional File 4). The reported resolutions are summarized as lateral (L) and axial (A) resolutions for electron microscopy (EM), X-ray microscopy (X-ray), and light microscopy (LM) methods in Figure 3B. Each dataset is plotted as a separate datapoint in Figure 3C. Overall, 72 datasets (61%) were generated using EM techniques, 29 (24.6%) with X-ray techniques, and 17 (14.4%) with optical techniques. As expected, the highest resolution was achieved by electron microscopy modalities (ET, FIB-SEM, SBF-SEM, ssTEM, STEM, FE-SEM, SEM-AT), followed by X-ray based methodologies (SXT, HXT, X-ray diffraction, STXM), and the lowest and most variable resolution in optical data (fluorescence microscopy, optical tomography). Isotropic resolution, which is highly advantageous for structural accuracy and morphological analyses (Chen et al., 2024; Le Gros et al., 2005), was far more common in X-ray based techniques (26/27 datasets, 92.9%, counting only those reporting both axial and lateral resolution), than in EM-based (25/56, 53.6%) or light-based (0/16) modalities.

Imaging resolution is therefore unique for each study, chosen within the available range to optimize feasibility and information density for the particular question or goal, and is a critical methodological parameter in methods reporting for critical interpretation of results, reproducibility, and evidence synthesis. In extracting the reported resolution from each study, we found that imaging resolution was not clearly reported for 18 datasets from 14 studies, in some cases inconsistently stated (conflicting and/or ambiguous mentions throughout the text), while in others, resolution was incompletely reported (e.g. axial (z), but not lateral (xy), resolution clearly stated, Figure 3C, right panel).

In summary, our results show that many modalities are capable of producing whole-cell image data, but X-ray based techniques, particularly soft X-ray tomography, have so far been most successfully leveraged to produce high resolution, isotropic, high-throughput whole cell datasets.

#### Cell type and sample size

The potential to pool and compare findings across studies depends largely on the biological similarity of the data, in particular, the cell type or model organism imaged. The same cell type imaged in multiple studies (with varying sample sizes) offers the potential to cross-reference and pool data across modalities and experimental conditions. The range of cell types represented across the corpus will point to appropriate groupings and comparisons for cross-cell type syntheses.

A wide range of cell types are covered in the corpus, including yeast, protists, plants, invertebrate animals, and both cultured and *in situ* mammalian cell types from rodents, non-human primates, and humans (Fig. 4A). In total, we found 80 distinct cell types across 118 datasets. The most strongly-represented cell types (in terms number of separate datasets and number of cells imaged in total across datasets, which we call “data density”), are highlighted in Table 1.

**Figure 4.**
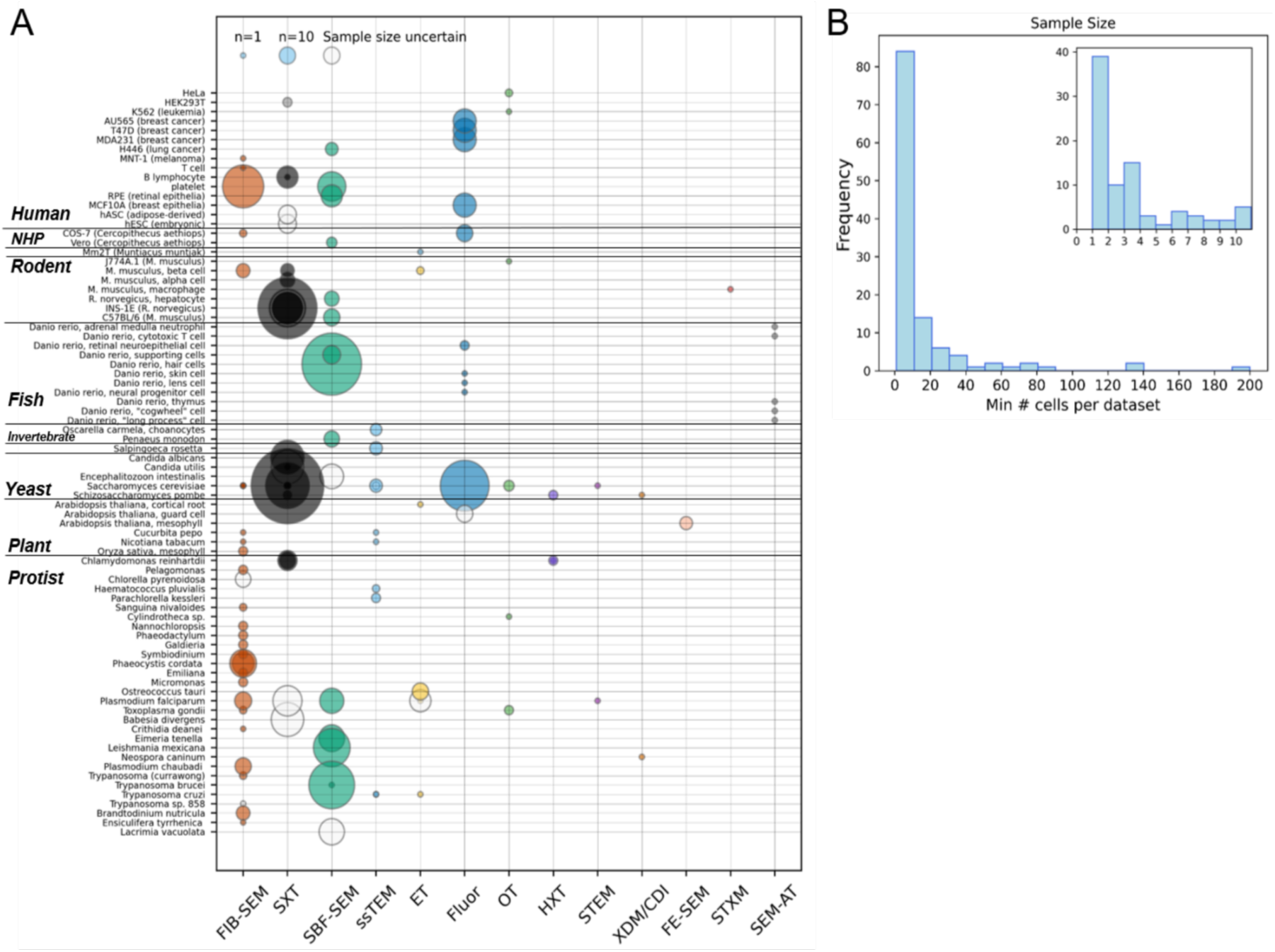
A) Datasets summarized according to cell type and modality. Each bubble represents one dataset, and bubble size represents reported sample size (refer to legend, top left), colored according to modality for clearly-reported sample size, and white for minimum known sample size when not clearly reported. B) Distribution of sample sizes, according to minimum known sample size. Inset show distribution for studies with sample sizes <=10.

**Table 1.**
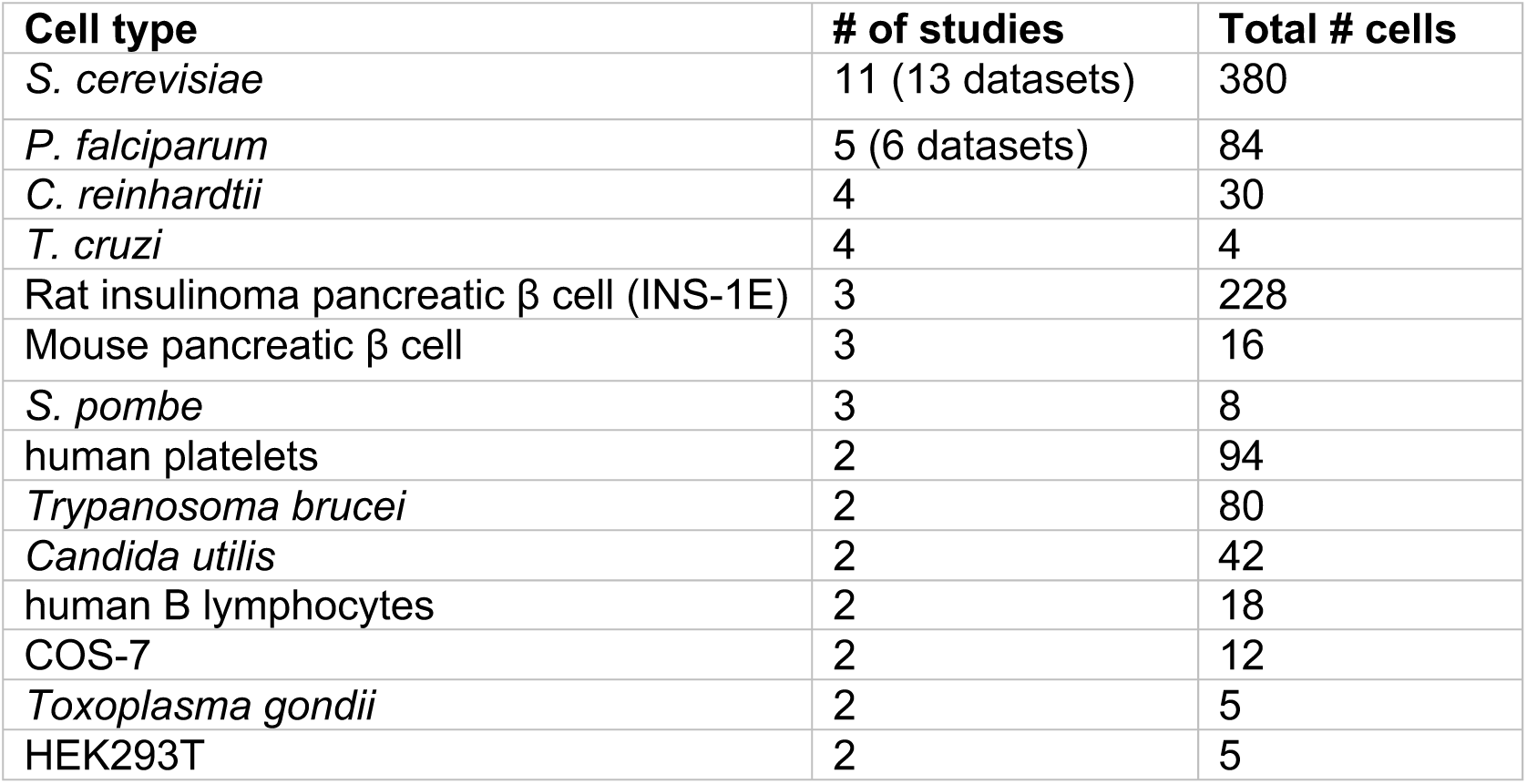
Cell types with highest data density. A summary of data available for cell types/species recurring in multiple studies across the corpus.

| Cell type | # of studies | Total # cells |
| --- | --- | --- |
| *S. cerevisiae* | 11 (13 datasets) | 380 |
| *P. falciparum* | 5 (6 datasets) | 84 |
| *C. reinhardtii* | 4 | 30 |
| *T. cruzi* | 4 | 4 |
| Rat insulinoma pancreatic $\beta$ cell (INS-1E) | 3 | 228 |
| Mouse pancreatic $\beta$ cell | 3 | 16 |
| *S. pombe* | 3 | 8 |
| human platelets | 2 | 94 |
| *Trypanosoma brucei* | 2 | 80 |
| *Candida utilis* | 2 | 42 |
| human B lymphocytes | 2 | 18 |
| COS-7 | 2 | 12 |
| *Toxoplasma gondii* | 2 | 5 |
| HEK293T | 2 | 5 |

Most studies had small sample sizes, defined as the number of cells fully reconstructed in 3D with multiple organelle volumetric information compatible with our inclusion criteria, with 81 (68.6%) of datasets reporting n<=10 whole cell images, and 36 (30.5%) studies consisting of a single cell (Figure 4B). Single-cell datasets were more common in earlier papers, and frequently occurred in technical papers – 16 of the 89 studies were technical papers reporting a single whole-cell reconstruction, representing half (51.6%) of the 31 technical studies and 66.7% of the single-cell studies. This suggests that the first ten or so years of our dataset (∼2005-2015) could be described as technical early days, but the advances during this time paved the way for more biologically-focused research studies with larger datasets in the following decade (∼2015-today). Sample sizes were largest in SXT (up to n=200) and SBF-SEM (up to n=141) studies. In 14 datasets, the study sample size, was either reported inconsistently or incompletely, in which case we reported a minimum known sample size, or else was not reported at all, resulting in missing data in our analysis (Figure 4A, white bubbles).

### Organelles and anatomical completeness

Our primary inclusion criterion was the presence of at least two organelles within the same whole-cell reconstruction. We argue that for visualizing cell anatomy, at least two different structures must be present in order to provide spatial context for one another. Furthermore, as pairwise spatial and morphological inter-organelle interactions are building blocks of the whole-cell organelle network, the presence of two organelles reveals at least one pair whose interaction can be described. We therefore extracted the list of organelles reconstructed in each dataset to assess the potential for analysis at the single-organelle, inter-organelle, and cell anatomical levels.

Figure 5A shows the cumulative organelle data per cell type across the corpus. The nucleus and mitochondria are the most commonly segmented organelles across all datasets (106 and 91 instances respectively, accounting for multiple distinct cell types per paper in some cases), followed by vacuoles/lysosomes (53 instances) endoplasmic reticulum (28 instances), lipid droplets (27 instances), Golgi (26 instances), chloroplast/plastid (27 instances), vesicles (18 instances), flagella/cilia (17 instances), peroxisome (7 instances), centrosome/basal body (5 instances), endosome (5 instances), multivesicular bodies (1 instance), and autophagosome (1 instance). Some organisms may appear to have only one or no organelles included despite our inclusion criterion requiring imaging of two or more organelles. This is due to the fact that the heatmap only includes “common organelles” present in all or most eukaryotes. The blank lines represent studies which image and reconstruct only specialized or cell type-specific organelles, such as the specialized secretory organelles (microneme, rhoptry, dense granules) or conoid structures in *Toxoplasma gondii* (Koutsogiannis Z *et al*., 2023; Paredes-Santos TC *et al*., 2012).

**Figure 5.**
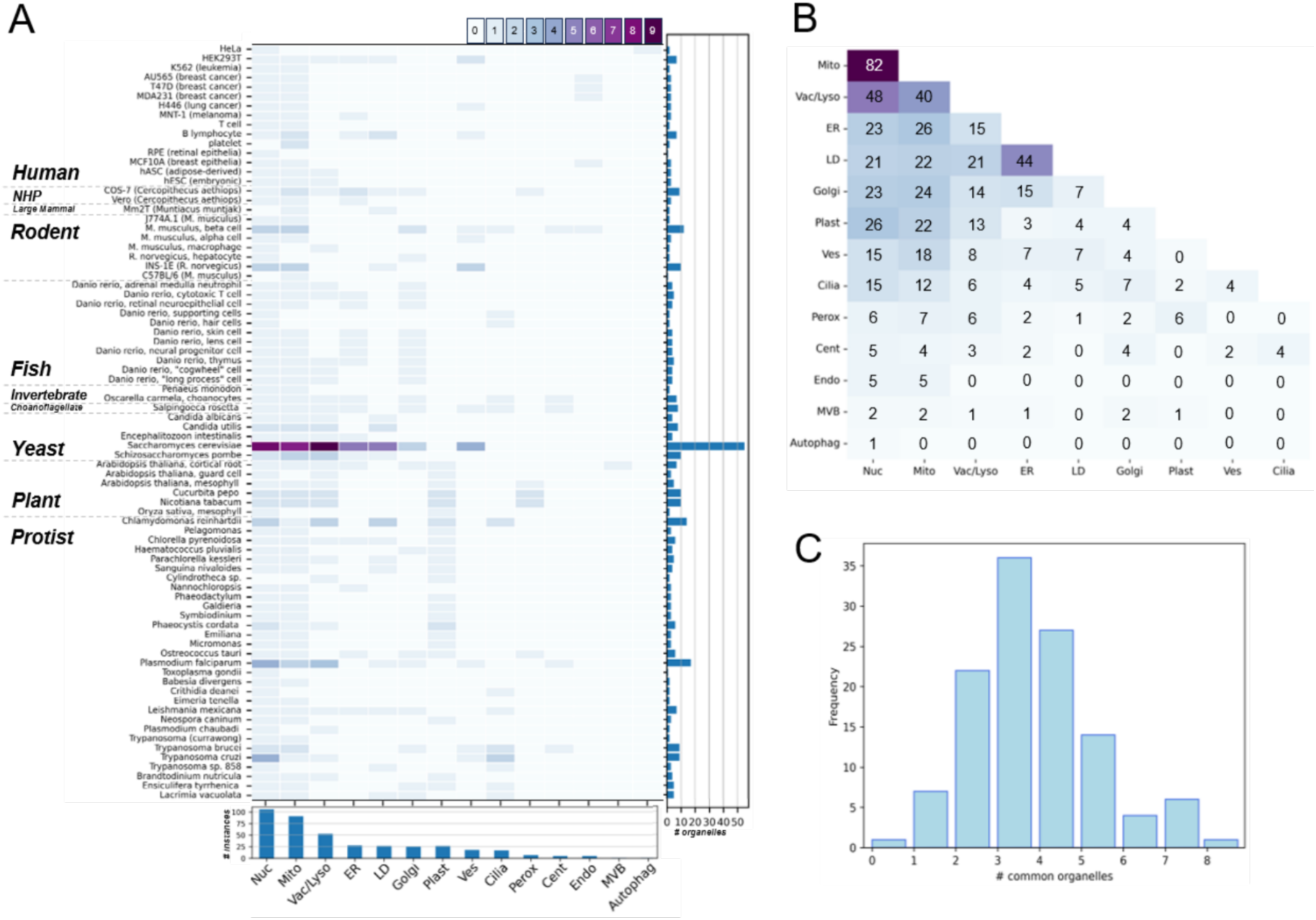
Representation of common organelles across datasets. A) Organelle data collected for each cell type represented across 118 datasets, excluding specialized or cell type-specific organelles. Heatmap color represents the number of independent datasets in which the organelle appears. Summary bar graphs show the total number of organelle instances per cell type (right) and total number of instances per organelle across cell types (bottom) (Nuc – nucleus, Mito – mitochondria, Vac/Lyso – vacuole/lysosome, ER – endoplasmic reticulum, LD – lipid droplet, Golgi – Golgi body, Plast – plastid/chloroplast, Ves – vesicle, Cilia – cilia/flagella, Perox – peroxisome, Cent – centrosome/centriole/basal body, Endo – endosome, MVB – multivesicular body, Autophag – autophagosome.) B) Organelle combinations. Heatmap represents the number of instances of each organelle pair co-imaged in the same dataset. C) Distribution of anatomical completeness, defined by the number of distinct common organelles coinciding in a single dataset.

In some cases, we find one common organelle in combination with specialized organelles, such as the study in C57BL/6 resting mouse platelets (mitochondria, dense granules, alpha-granules, canalicular system) (Pokrovskaya I *et al*., 2021). The recurrence of cell types in multiple studies enables potential examination of organelle morphologies across datasets, experimental designs, and biological contexts. While not all studies imaging the same cell type segment the same set of organelles, the most frequently-imaged specific organelles are found among the best-represented cell types: *S. cerevisiae* vacuoles (12 instances), nuclei (11), mitochondria (10), ER (7),lipid droplets (7), and vesicles (5); *P. falciparum* nuclei (5) and vacuoles (4), and *T. cruzi* nuclei (5).

The rationale for requiring at least two organelles per segmented cell in the dataset is 1) the supposition that a single internal landmark is insufficient to constitute an anatomical pattern, and 2) to reveal spatial interactions between organelles in the 3D context of the entire cell. Overall, we found 59 pairwise organelle combinations occurring across the corpus, of which 35 had at least five independent instances, suggesting rich potential for comparison and synthesis of each organelle-organelle relationship across many cell types and contexts (Figure 5B). The most common organelle co-occurrences are mitochondria-nucleus (82 instances across 65 cell types), nucleus-vacuole/lysosome (48 instances across 36 cell types), and mitochondria-vacuole/lysosome (40 instances across 33 cell types). These cases will enable within-cell type synthesis of similar data (e.g. analysis of a specific organelle pair, as well as cross-study comparison, which can potentially isolate biological and technical differences.

The ultimate requirement for an understanding of cell anatomy and modeling whole-cell patterns and inter-organelle interaction networks is to be able to visualize as many organelles simultaneously as possible. The number of “common organelles” per reconstruction, termed “anatomical completeness” is distributed with a peak at 3-4 organelles (Figure 5C). The most anatomically complete reconstructions are in choanoflagellate *S. rosetta* (Laundon et al., 2019) with 8 organelles, and 7 organelles in *L. mexicana* (Hair et al., 2024), sponge *O. carmella* choanocyte (Laundon et al., 2019), *A. thaliana* cortical root cell (Cui Y et al., 2019), and *S. cerevisiae* (Wei et al., 2012).

### Morphological metrics and Quantitative analyses

The unique advantage of high-resolution 3D structural data of whole cells is their capacity to provide rich and precise quantitative morphological and spatial information. Future meta-analysis of organelle features relies on the presence of comparable quantitative measurements across studies. An assessment of quantitative analyses performed across studies revealed a limited set of organelle geometric features measured (Figure 6A). Volume was the most frequently measured property (65 studies, 73%), followed by volume fraction (organelle-to-cell volume ratio, 50 studies, 56%). Also common were surface area (34 studies, 38%), count/copy number (29 studies, 32.6%) and linear dimensions such as length and diameter (27 studies, 30%). Less frequent, and more variable, were measures of density such as LAC or electron density and positioning/distance (n=18 each, 20.2%), organelle contacts/associations and shape (n=14 each, 15.7%). Organelle volume scaling trends were calculated in 7 studies (7.9%) and three studies measured refractive index. One study devised a method to quantify subcellular crowding of organelles (“congestion index”, Loconte V *et al*., 2022). These frequencies roughly reflect the general ease of measurement and conceptual familiarity of these metrics - volume and surface area are generally trivial to measure, albeit with variable accuracy, with many standard software tools automatically generating these calculations. Count and dimensions can be straightforwardly manually or semi-automatically calculated. Distance and contact frequency, shape, and crowding require more complex calculations and computational operations, and can be interpreted in variable ways. A sizable fraction of studies (n=14, 15.7%) performed no quantitative analysis.

**Figure 6.**
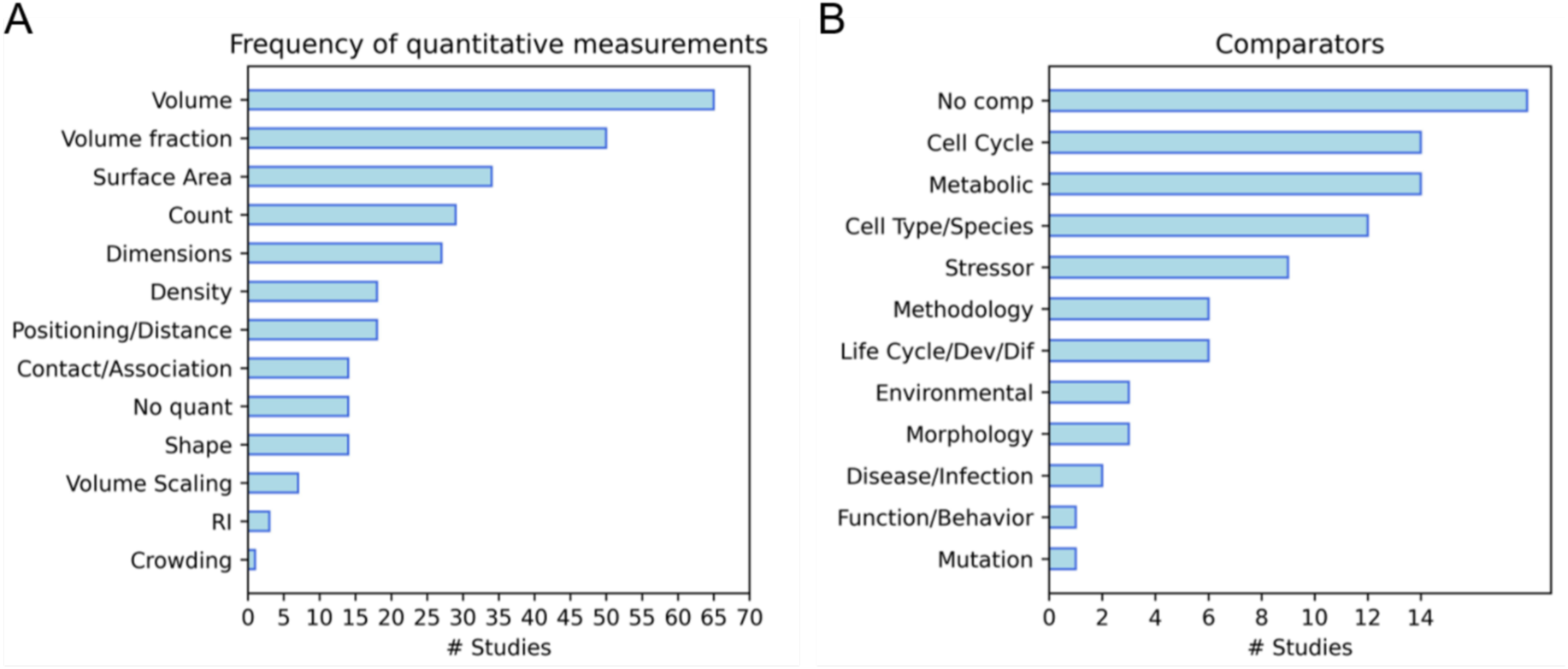
A) Quantitative analyses performed on whole-cell reconstruction data. Dimensions, density, positioning/distance, contact/association, and shape are all groupings including different measurements conveying similar properties. (No quant – studies with no quantitative analysis; RI – refractive index) B) Frequency of comparator classes by study. Within-study comparable conditions or perturbations were grouped into general classes. (“No comp” – studies with no comparator; Life Cycle/Dev/Dif – stages of life cycle, development, or differentiation)

### Comparators

While whole-cell structural data is largely static, cellular structure is in fact highly dynamic and variable across cells. We recorded the experimentally compared conditions between samples (“comparators”), reasoning that variety in conditions for a single cell type could suggest functional organelle and anatomical variability, while overlap in conditions across cell types could reveal generalizable context-specific organelle and anatomical trends. Most studies (71, 79.8%) included comparator data across at least one cellular perturbation or condition, which we grouped into broad classes (Figure 6B). The most common of these are cell cycle stages and metabolic conditions, such as glucose stimulation of pancreatic cells (14 studies each). Other comparators, in decreasing order of frequency, were cross-cell type comparisons (n=12), stressors such as drug treatments (n=9), methodology, such as fixation methods or imaging modalities (n=6), stages of life cycle, development, or differentiation (n=6), environmental conditions, such as light exposure (n=3), morphology, such as shape or size (n=3), disease or infection states (n=2), functional or behavioral states, such as cell-cell communication (n=1), and genetic mutation (n=1). The predominance of cell cycle stages and cell type as the primary comparators highlights the potential of whole-cell imaging to serve as a basis for defining cell type and functional cell state-related cellular reorganizations.

In summary, the whole-cell multi-organelle imaging corpus includes 89 published studies between 2004 and 2024 from over 11,000 initial PubMed results. In total, the dataset represents the whole-cell imaging and reconstruction of at least 1690 individual cells, of 80 eukaryotic cell types from protists to humans, across 118 independent datasets, 13 imaging modalities, at least 13 common organelles in many different combinations, 71 perturbative or comparative conditions, and at least 12 types of morphological metrics quantitatively analyzed.

## LIMITATIONS

Our scoping study is, to our knowledge, one of the first applications of scoping review methodology to the field of whole-cell microscopy, and to fundamental cell biology more broadly. Our method comes with limitations arising from the iterative nature of proof-of-concept methods development, the breadth and heterogeneity of the literature corpus, and the size of our team. One of the goals of a formal systematic literature synthesis is to be as comprehensive as possible, spanning peer-reviewed and grey literature sources as thoroughly as possible. Our approach focused specifically on PubMed, but ideally, other databases such as Web of Science and EMBASE, would be included as well. Although our PubMed search was extremely broad relative to our final inclusion set, which comprised only less than 1% of the original search results, we cannot be sure how many relevant studies might have been missed by the search entirely. Furthermore, grey literature sources such as dissertations and online image data hosting sites without corresponding publications would be of relevance to a comprehensive set of cell anatomy data, but would be missed by our strategy. For example, the Allen Institute for Cell Science hosts a collection of 3D confocal images in human iPSCs, a subset of which would fit our inclusion criteria, but these datasets have not been directly thoroughly described in a publication, despite their inclusion within a large dataset underlying many impactful publications (*Cell Line Catalog*, 2024). Furthermore, as of January 2025, our search string retrieves 300 new results since our last round of curation in May 2024. The dataset and analysis presented here should be seen only as an initial characterization which will be continually updated as the literature base expands. We intend to establish our growing curated corpus and analysis as a publicly available, living database of cell anatomy data and field trends, inviting community input to identify gaps and curate the corpus as thoroughly as possible.

Another limitation of our approach, as of any systematic research synthesis, is that the choice of strict inclusion criteria necessarily excludes potentially valuable studies. These boundary cases help to define the field of whole-cell reconstructions and reveal peripheral sources of knowledge that can be incorporated into follow-up studies depending on the specific biological question. One core criterion was the need to define “*whole cell*” strictly, requiring that the full cell periphery is clearly delineated. The rationale for this is that, first, the full anatomy of the cell requires that no part of the cell is missing from the image, and second, the intact cell boundary is needed for accurate measurement of cell volume, which we consider to be the critical common measurement required for any cross-study quantitative analysis as the denominator for organelle volume ratio. Therefore, many studies reporting mostly complete cell volumes, in some cases even describing the data as “whole cell” despite missing small cell edges, had to be excluded. This was often the case for adherent cultured cells, which spread out flat on culture dishes with thin, faint, irregular protrusions when sparsely plated, and for very large, irregularly shaped cell types such as muscle cells and neurons, which are extremely difficult to perfectly capture in their entirety *in situ* or in culture. In other cases, we found studies of tissue sections containing partial reconstructions of multiple cells clipped at the section boundary, including organelle volumes and volume ratios pooled and normalized across the entire section volume, but no single cell fully reconstructed. Therefore, our strict “whole cell” criterion introduces a cell type bias into our corpus. Moving forward, a case could be justified for softening this criterion to estimate rather than directly calculate cell volume. The systematic nature of our screening process means that steps can be back-tracked to retrieve specific studies previously excluded for either of these reasons and include them under an adjusted set of criteria.

## DISCUSSION

We have created a comprehensive corpus of whole-cell imaging literature spanning a wide breadth of modalities, cell types, organelles, and contexts through a novel application of formal scoping review methodology. There is substantial comparability across studies, highlighting opportunities for further meta-scientific and synthetic studies building on this corpus. Future directions include systematic review and meta-analysis approaches to understand how organelles typically self-organize and interact within the whole cell space across biological contexts, including qualitative categorizations of cell anatomy, and meta-analyses of key morphometric measurements.

Although our search is not exhaustive, this corpus comprises a foundational evidence base revealing an emerging field of significant, and rapidly increasing, volume and scholarly interconnectedness, unified by a specific data type – whole cell, multi-organelle 3D reconstruction (Figures 1 and S1). Our systematic curation and meta-scientific characterization of these remarkably information-dense structural data, from raw data to processed images to derived measurements, enables a range future secondary analysis and synthesis efforts. The substantial methodological and biological overlap opens a wide range of possibilities for within- and cross-group synthetic studies to uncover novel insights that would otherwise be difficult to access through traditional primary research methods. Data in a specific cell type pooled across datasets, modalities, and conditions provide opportunities to query the consistency of similar metrics across studies, integrate disparate findings into an organism-centric model, and compare distinct imaging modalities and parameters applied to the same experimental system. Summarizing quantitative analyses and extending them where possible allows for a more comprehensive quantitative understanding of generalizable principles of organization, an assessment of single organelle, between-organelle, and cell-wide feature variability across contexts, and provide a strong empirical foundation for integrative modeling efforts (Loconte et al., 2023). Publicly shared original data enable novel analyses and the development of new data integration and secondary analysis workflows. Ultimately, work building on our corpus promises to reveal emergent patterns of cell anatomy across cell types and contexts, and insights into the constraints and variations embedded in these patterns as a result of cellular geometry and organelle interactions.

We intend for our dataset and analysis to become a growing tool for the scientific community. In addition to generalized, high level synthesis, the corpus can be subdivided in many ways to enable inquiry into specific cell types, organelles and organelle interactions, biological perturbations, imaging and image analysis approaches. In the near future, we intend to make the corpus easily accessible, interactive, and kept up-to-date so that researchers interested in any component of the corpus can easily contribute to and draw upon it for their own investigations. We hope this work will galvanize whole-cell imaging and analysis researchers, synthesis researchers, and anyone else interested in organelle-to-whole cell mesoscale structural biology to create a community, collaborating to refine the ongoing curation and re-use of whole-cell data, refine and unify the generation and reporting of novel work, and engaging in discourse around terminology, best practices, optimal applications for existing imaging technologies and needs for technical advancement, and a broad vision of the future of the whole-cell imaging and cell anatomy field.

The range of imaging modalities and technical parameters represented across our corpus demonstrates the versatility of approaches available to achieve whole-cell anatomy data. It also provides opportunities to directly compare these approaches and trends in how they are applied, identify common challenges, and motivate recommendations for future work. We find that the current whole-cell imaging landscape, on the whole, is biased toward small cell types and small sample sizes, with a single cell reconstruction reported in nearly one-third of studies. This is largely due to the time-, cost-, and labor-intensive nature of whole-cell image acquisition, reconstruction and multi-organelle 3D segmentation, particularly at sub-100nm resolution. The 13 imaging modalities represented each offer particular advantages and disadvantages, with unique tradeoffs between resolution, throughput, native structure preservation, and overall tractability depending on the application. For example, the two dominant modalities, SXT and FIB-SEM, are both capable of achieving high isotropic resolution with some degree of quantitative throughput. SXT generally achieves isotropic 30-60nm resolution with minimal sample damage or tomographic artifacts, and is capable of producing throughput in the hundreds per study. However, cell size is strictly capped by the ∼10um diameter of the capillary used to hold the samples, limiting the cell types one can study by this method, and access is limited by the requirement for beamline time. FIB-SEM excels for very high-resolution reconstructions (∼5 nm), but is limited by throughput and, as a sample-destructive method, is more likely to introduce ultrastructural artifacts. Extensive reviews of strengths and weaknesses across volumetric imaging modalities have been discussed in depth elsewhere (Hanssen et al., 2012; Loconte et al., 2021; Loconte V et al., 2022), and technical capabilities for all modalities are constantly being improved.

However, the limiting factor for maximizing the utility of whole-cell image data is perhaps not data quality or abundance itself, but rather the limited use and reuse of these incredibly rich datasets, potentially leaving much of the embedded information untapped. We found that the publication of whole-cell imaging studies is accelerating, with growing trends toward open-access publication and data sharing, suggesting good current and future potential for meta-scientific studies and secondary re-analysis of datasets (Fig. 2). However, only a small subset of our corpus made original data publicly available, and we found inconsistent reporting of key methodological details, such as imaging resolution and sample size. Such metadata items are crucial for meta-scientific analysis, including secondary analysis and synthesis research as well as within-study reproducibility and cross-study replicability assessment. Currently, there is increasing momentum in the development of open science and data sharing guidelines and implementation led by international collaborations of scientists (FAIR, DORA), funding bodies (NIH, NSF), and publishers. Many data type-specific metadata standards have been developed for bioimage data by community consensus (Global BioImaging, REMBI, foundingGIDE, Open Microscopy Environment) and the growth of image dataset repositories (EMBD, EMPIAR, IDR, SSBD, NIH Imaging Data Commons). Image data, particularly high-content volumetric imaging such as whole-cell reconstruction datasets, present particular challenges for data sharing, such as the large size and complexity of files and datasets, irregular file formats. Ongoing efforts to develop novel bioimage file formats (OME-Zarr (Moore et al., 2023); OME-NGFF (Moore et al., 2021)) and accessibility tools (BioFileFinder) promise to help tackle some of these barriers, and lead to new solutions as the public image data ecosystem scales up. There may be factors of particular interest to the emerging scientific community generating and using whole-cell imaging data with regards to specific imaging modalities and data types, such as segmentation and 3D reconstruction parameters. As the publication of whole-cell imaging studies accelerates, we envision that community discourse and establishment of specific standards for methodological design, reporting, and data sharing could be of broad benefit. We hope to see a coordinated effort to develop open science practices to maximize the utility of these costly-to-produce data.

While interest in open science and data-reuse is quite mainstream and rapidly growing in the sphere of cell and molecular biology, the range of approaches currently used are largely focused on big -omics data informatics. The past two decades have yielded the independent development of synthesis and integration methods in some specific contexts, such as GWAS meta-analyses in genomics, computationally-assembled gene- and protein interaction networks in bioinformatics, and massive consortium collaborations to map cell types and beyond (“Meta-Analysis in Basic Biology,” 2016). There is still a clear need for robust knowledge and data synthesis in biology, driven by widely recognized challenges associated with “data deluges” (“The Data Deluge,” 2012; Zhang, 2024) and the exponential growth of scientific output (Hanson et al., 2024; Pautasso, 2012). The monumental goal of constructing meaningful holistic understandings of highly complex biological systems by merging a rich tradition of narrowly focused mechanistic studies and molecular parts lists, with interdisciplinary systems, biophysical, evolutionary, synthetic, and engineering perspectives (Rafelski & Theriot, 2024), requires robust frameworks to comprehensively and rigorously synthesize data, knowledge, and interpretations across disciplinary boundaries, aided by a high-level understanding of field trends revealed by meta-scientific analyses.

There is at present limited familiarity in the fundamental biology field of evidence synthesis methods widely used in many disciplines including the clinical and social sciences. These are extensive, carefully developed toolkits supported by synthesis researchers and large international collaborations and initiatives such as Cochrane and the Joanna Briggs Institute. Systematic research syntheses in fundamental cell biology, particularly intermediate-scale syntheses to address specific biological questions or hypotheses, are beginning to come on to the scene in recent years, still rare and methodologically disparate (Gillooly et al., 2015; Opalek et al., 2023; Rutkowska et al., 2020; Sercel et al., 2023; Trumpff et al., 2021).

The present work illustrates the unique considerations, challenges, and scientific and meta-scientific potential of developing an evidence synthesis toolkit tailored to fundamental cell biology. In contrast to translational and applied disciplines, particularly those based on animal or human subjects, exploratory fields such as fundamental cell biology marked by high heterogeneity in their technical repertoire, study design, experimental and reporting practices, philosophical and sociological standards for epistemology and argument construction, and conceptual and linguistic frameworks. This heterogeneity presents unique and significant challenges for the application of highly standardized, guideline-driven evidence synthesis methods. For example, development of a narrow and specific bibliographic database search strategy is particularly challenging. Firstly, deep and broad technical expertise is required to understand the scope of research likely to contain a particular data type, specific experiment, or conceptual theme. Second, highly varied terminology used across the literature, and particularly in the lack of concrete norms for titles and abstracts results in a tendency toward very large search result corpuses, preferring comprehensiveness over specificity. The search scope and resulting corpus may therefore be larger than is the accepted norm in other disciplines. Therefore, inclusion criteria must be carefully justified and iteratively refined, and robust stepwise methods for screening and data extraction, ideally carried out by teams of 2-4 expert investigators working in parallel to minimize bias, are needed. In general, a typical systematic review process can take 1-1.5 years of painstaking manual work conducted and validated by a dedicated team.

The prospect of employing AI tools to aid in these intensive steps is attractive, and being actively explored by many scientific groups and software companies (Bolaños et al., 2024; Yip et al., 2024). However, due to the goal of maximum comprehensiveness and need for a careful balance between rigor and pragmatic flexibility to handle the highly heterogeneous and nuanced nature of the fundamental biology literature, the requirement for human judgement and validation is unlikely to be replaced by AI in the immediate future. We could envision a manually curated scoping study, such as this one, forming the foundation for a novel synthesis effort, and in tandem, creating a robust training dataset for an AI synthesis assistant, while further refinement, establishment of a living corpus, and updating with novel publications as they are released, can be accomplished with the aid of the trained AI model and community curation and validation. We look forward to a bright future in which specialized synthesis researchers, in collaboration with the expert community and automated tools, help to create a thriving ecosystem for novel discovery and meta-research to derive the maximal benefit from the immense volume and richness of biological literature and data.

## METHODS

### Search strategy

We developed our search strategy starting with an initial set of 11 target papers (Decelle J et al., 2021; Heinrich L et al., 2021; Henderson et al., 2007; Laundon et al., 2019; Müller et al., 2021; Uchida et al., 2009, 2011; Uwizeye et al., 2021a; Valm et al., 2017; Wei et al., 2012; White et al., 2020)], thoroughly combed for cell attribute-(e.g. “cell organization”, “subcellular architecture”), mesoscale-, and imaging-related terms. These terms were used to devise an iterative series of search strings for PubMed aimed at retrieving a defined corpus that includes the full initial set. While an initial attempt focused on cell anatomy-related terms was unsuccessful due to the heterogeneity and inconsistent usage of such conceptual language, iterations on search strategies consisting of 3D imaging and mesoscale (cell/subcellular and organelle-scale) terms successfully retrieved the initial papers.

> ((“Imaging, Three-Dimensional“[MH] OR “3D imaging“[tiab] OR “three-dimensional imaging“[tiab] OR “3-D imaging“[tiab] OR “Soft X-ray tomography“[tiab] OR (3-D[tiab] OR 3D[tiab] OR “three dimensional“[tiab] OR “three-dimensional“[tiab]) AND (tomography[tiab] OR microscopy[tiab] OR Microscopy[MH] OR Tomography[MH] OR imaging[tiab]))
>
> AND
>
> ((ultrastructure[MeSH Subheading]) OR (Organelles[mh] OR organell*[tiab]) OR (intracell*[tiab] OR intra-cell*[tiab]) OR (subcell*[tiab] OR sub-cell*[tiab]) OR (((whole[tiab] OR entire[tiab]) AND cell[tiab]) OR “whole-cell“[tiab] or “whole cell” OR “entire cell“[tiab]))) NOT Review[pt]

We set our search to begin at 2005 due to the emergence of technological capability to perform whole-cell microscopy and reconstructions around this time (Le Gros et al., 2005). However, this missed one key earlier study, which was added *post hoc* (Larabell & Le Gros, 2004). The most recent PubMed search update was performed on May 23, 2024.

### Screening

Our search strategy returned over 11000 results from PubMed. Because our search was aimed at a particular data type (whole-cell volumetric microscopy) rather than a specific research topic, our target corpus is highly heterogeneous, spanning a wide range of publication venues, key terminology, and other bibliometric features. Therefore, our primary objective was to retrieve our initial set as well as any studies with even a marginal potential of containing whole-cell microscopy data, yielding a highly sensitive but not specific search.

To curate our dataset, we performed three rounds of screening according to the following strictly-defined inclusion criteria:

1. Cell anatomy is necessarily a whole-cell property. The study must have collected original three-dimensional *whole-cell* images in eukaryotic systems, described in the primary study in the form of direct visualizations (still images and/or supplementary movies displaying volumetric rotations), and/or quantitative analyses necessarily taken from whole-cell data (e.g. organelle-to-cell volume ratios). “Whole-cell” requires that the entire cellular boundary (cell membrane or cell wall, or other marker showing the continuous cell periphery) is captured within the image, such that the cellular volume can be calculated.
2. Volumetric information for at least two *organelles* concurrently must be included in the whole-cell volume. We reasoned that at least two subcellular structures must be visible in relation to one another for the concept of “anatomy” to be applicable. This criterion requires a full volumetric reconstruction of the multi-organelle cell, with organelle identities clearly annotated (typically in the form of a segmentation). The reconstructed data should either be directly shown as an image in figures, movie files, or associated data, or necessarily generated to produce published analysis, such as volume calculations. Although the usage of the term “organelle” has expanded in recent years, we used a traditional definition, referring to a structurally and functionally distinct, membrane-bound, intracellular compartment.

Curating a set of target papers from this large set of search results required a series of increasingly thorough screening steps – title and PubMed summary, abstract, and full text – conducted by a team of three reviewers. Search results and screening decisions for each stage, including exclusion reasons, along with information extracted from initial target papers are provided in Supporting File 1.

#### Title/summary

The first screening step was a single-reviewer rapid screen through search result pages in the web browser (Mozilla Firefox) of titles and short three-line PubMed-generated summary statements. Studies that were obviously irrelevant, focused on the wrong biological scale, or purely technical with no biological data were excluded. Studies that clearly described whole-cell imaging and/or structures of entire cells were included. If unclear, a skim of the abstract (and PMC figures, if available) was done, and studies that were still ambiguous were included for further screening. This step was performed by one reviewer (MM), and excluded 95% of results.

#### Abstract

A custom Python (v3) script using BioEntrez module of Biopython (Cock et al., 2009) was used to pull abstracts from PubMed e-utilities (*Entrez Programming Utilities Help*, 2010) and record screening decisions from one reviewer. Studies were either included, excluded with exclusion reasoning recorded, or marked for full text screening. A 10% subset was screened in parallel by a second reviewer (WM) to check inter-rater consistency. A low level of consistency was initially found due to differing definitions of “organelle”. This led to a refinement of our working definition of “organelle” for the purpose of screening criteria (see above), and the full set of abstract screening was performed in parallel by two independent reviewers (MM & BW). This step yielded 60 clear hits (10.8%), excluded 184 (32.3%), and sequestered 326 studies for full text review.

#### Full text screening

Full texts, including supplementary data, were screened in parallel by two reviewers (MM & BW). Screening decisions were recorded in Covidence software (Veritas Health Innovation, n.d.), along with reasons for exclusion. Exclusion reasons were categorized as: missing 3D data, including cases where 3D data is textually described but not included as data in the paper (“Not 3D”); imaging only at subcellular or multicellular/tissue scales (“Wrong scale”); cells only partially imaged and/or reconstructed, or with cut-off edges such that cell volume cannot be calculated (“Not whole-cell”); raw data missing organelle segmentation (“Not segmentation”); fewer than 2 organelles segmented (“Not multi-organelle”); data reused from other studies (“No original data”); review or perspective articles (“Wrong article type”); and a general lack of whole-cell, multi-organelle reconstruction data (“No relevant data”).

We also designated 39 studies as “Borderlines” if they represented edge cases with regards to our inclusion criteria, but nevertheless contained relevant data, analysis, or insights, therefore not warranting complete exclusion from our dataset, summarized in Additional File 1. One borderline category is “whole-cell data not shown”, including studies in which the target data type is implied or explicitly described in the text but not included in figures or associated data files, or provided in supplement where files are inaccessible. This includes several studies that partially reconstructed prohibitively large cell types (Trebichalská et al., 2021), as well as reconstructed tissue volumes from which while relevant metrics are quantified, such as organelle volume/volumetric ratios, calculated across the tissue block rather than in individual cells, as distinct cell boundaries could not be delineated. In some cases, the whole-cell images had slightly clipped cell edges due to nuances such as cells in cultures spreading their edges and protrusions far from the main cell body on a culture substrate, or an elongated cell type such as myocytes. We chose to exclude such examples but include others such as images of protists with flagella partially cut-off, due to the fact a missing edge of the cell body leaves key morphometrics, such as cell volume and shape, impossible to calculate, and creates uncertainties about holistic cell anatomy due missing information, whereas missing part of a cilium or flagellum would only affect analyses of ciliary length, but would preserve the ability to calculate organelle to cell size ratios. Other borderline categories include: organelle identities uncertain; ineligible structures defined as “organelles“; a very minor volume of the cell edge is cut off in the image, technically violating the “whole cell” criterion; cell membrane not clearly delineated; the paper itself does not include the relevant data, but accompanies the establishment of a public repository or resource containing relevant data.

Disagreements in inclusion/exclusion decisions were discussed between reviewers, prompting slight adjustments to our protocol, such as establishing new exclusion reasons or borderline categories. One earlier study was added *post hoc* (Larabell & Le Gros 2004). Ultimately, 89 studies were fully included.

### Data Extraction and Analysis

We extracted information from studies to summarize study design, including imaging modality and resolution, cell types, sample sizes, organelles, comparative conditions, and quantitative analyses, only as pertains to information within the study directly describing or derived from whole-cell imaging data, as opposed to other types of data within the same paper. Data extraction was performed manually by two researchers independently (MM & BW), and subsequently discussed observations to reach consensus.

Data items were annotated on full text PDFs in Zotero (*Zotero | Your Personal Research Assistant*, n.d.) and entered into the data extraction template in Covidence (Veritas Health Innovation, n.d.), followed by inter-rater consensus discussions to resolve disagreements and inconsistencies. Extracted data were exported from Covidence as .csv, and manually processed and formatted in Excel. The full set of final data extraction items, including 89 included studies and 39 borderline studies, is provided in Additional File 2. Co-authorship networks were generated in VOSViewer (van Eck & Waltman, 2010) using PubMed files as input and the following settings: bibliographic data, all co-authorship, include documents with a large number of authors, allow unconnected items, and overlay visualization. Citation network was created in Litmaps (*Litmaps (Version 2025-01-16) [Search Tool].*, 2024) using the following settings: X axis: Publication Date, compact; Y axis: Map Connectivity; Size: Cite Count, linear interpolation; disable Position Editing, Show citation arrows, Avoid collisions. All other data analysis and plotting was performed in Python using Jupyter notebooks and the following packages: Matplotlib (Hunter, 2007), SciPy (Virtanen et al., 2020), Pandas (The pandas development team, 2020), Numpy (Harris et al., 2020), Seaborn (Waskom, 2021). Code and source data files are available at https://github.com/bdwucsf/Cell-Anatomy-Scoping-Review.

## Acknowledgements

This project was funded by the UCSF Program for Breakthrough Biomedical Research, funded in part by the Sandler Foundation, and NIH grant R35 GM130327. We owe a debt of gratitude to the UCSF librarians, Peggy Tahir, Josephine Tan, and Evans Whitaker, for evidence synthesis methods consultations and assistance with developing our search strategy. We thank Dr. Jane Maienschein, Dr. Susanne Rafelski, Dr. Sabina Leonelli, Dr. Matthew Akamatsu, Dr. Shane Jinson, Dr. Sam Church, and members of the Marshall lab for constructive feedback and support.

## ADDITIONAL FILES

**Additional File 1.** Step by step details of corpus construction, with each step of searching and screening listed in a separate tab. 0_Initial Target Papers: list of initial papers and extracted features exemplifying target article types, used to construct PubMed Search Strategy. 1_Pubmed Search (2005-5.28.24): Complete list of 11,104 PubMed search results. 2_Title Screening: PubMed metadata for 570 titles remaining after first screening step based on titles and PubMed summary statements. 3_Abstract Screening: Results of second screening step based on abstracts with PMID, inclusion decision (Include (I), Exclude (E), and Check Full Text (C) and brief justification for each study. 4a_Full Text Screening Includes: Metadata for 127 studies included after full text screening. 4b_Full Text Screening Excludes: Metadata for 256 studies excluded after full text screening. 5_Post hoc additions: Metadata for 1 study included after completion of bulk screening.

**Additional File 2.** Complete data extraction information for included studies after full text screening, with full includes (Data Extraction_Includes) and borderlines (Data Extraction_Borderlines) separated.

**Additional File 3.** Publicly available datasets. Parameters for 12 studies reporting public data availability, including dataset completeness, study design, links, amount and format of data.

**Additional File 4.** Complete information for 118 datasets. Each individual dataset from 89 studies is listed including specifics of imaging modality, resolution, cell type and taxonomy, sample size, and organelles.

**Additional File 5.** Data underlying analysis plotted in figures. Specific subsets of data used to plot figures is separated into tabs.

